# Visual network modularity and communication alterations in ADHD subtypes: evidence from source localized EEG and graph theoretical analysis

**DOI:** 10.1101/2024.05.16.594553

**Authors:** Amir Hossein Ghaderi, Shiva Taghizadeh, Mohammad Ali Nazari

## Abstract

The neurobiological basis of ADHD and its subtypes remains unclear, with inconsistent findings from studies using electrophysiology and neuroimaging. Some studies suggest ADHD-I is a distinct disorder, but there is also evidence of similar neural basis in ADHD-I and ADHD-C subtypes. This study investigates the neural basis of ADHD and its subtypes using a subnetwork modularity approach based on graph theoretical analysis of EEG data from 35 children aged 7-11. EEG was recorded in the eyes open condition and preprocessed. After preprocessing, data was analyzed using LORETA algorithm to estimate current densities in 84 regions of interest (ROIs) in the cortex and calculate functional connectivity between these ROIs in different EEG frequency bands. Then, we evaluated modularity of five functional brain networks (default mode, central control, salience, visual, and sensorimotor) using Newman modularity algorithm. Further, we evaluated edge betweenness centrality to assess communications between these functional brain networks. The study found that different brain networks have modularity in certain frequency bands, and ADHD groups showed reduced modularity of the visual network compared to normal groups in the alpha1 band (8-10 Hz). The communication between the visual network and other brain networks, except the salience network, was also reduced in ADHD groups (in the alpha1 band). However, there were no significant differences in the modularity of brain networks and communication among them between two ADHD subtypes. The results suggest a novel mechanism for ADHD involving lower intrinsic modularity in the visual network, disturbed communication between the visual network and other networks, and potential impact on the function of control and sensorimotor networks. Further, our results suggest that there may be a common neural basis for both subtypes, involving a shared disturbance in the modularity and connectivity of the ventral network. This supports the idea that ADHD-I and ADHD-C are subtypes within the same category and contradicts previous studies that suggest they are separate disorders.

## 1) Introduction

Attention Deficit Hyperactivity Disorder (ADHD), is a common developmental disorder that affects around 6% of children and young people globally (Faraone et al., 2021). According to the DSM-5, children with ADHD display symptoms of inattention and/or impulsivity-hyperactivity in at least two different environments, such as school and home. This condition is classified into three subtypes based on the severity of inattention or hyperactivity: inattentive (ADHD-I), hyperactive (ADHD-H), and combined (ADHD-C). However, despite this well-defined behavior-based classification, the neural basis of these subtypes is still unclear, and existing studies using electrophysiology or neuroimaging have not reached a consensus on the similarities or differences between ADHD subtypes (Aldemir et al., 2018; Clarke, Barry, McCarthy, & Selikowitz, 2001; Ercan et al., 2016; Fair et al., 2013; Amir Hossein Ghaderi, Nazari, Shahrokhi, & Darooneh, 2017; Grizenko, Paci, & Joober, 2009; Jacqueline F Saad et al., 2021).

Particularly, some studies have suggested that ADHD-I should be considered as a distinct disorder rather than a subtype of ADHD (Grizenko et al., 2009; Milich, Balentine, & Lynam, 2001; Z.-M. Wu et al., 2022). This notion has received both support and opposition from neurocognitive and neuroimaging assessments. In neurocognitive assessments, differences in visuospatial short-term and working memory between ADHD-I and ADHD-C has been highlighted in one study, suggesting more severe impairment in ADHD-C(Dovis, Van der Oord, Huizenga, Wiers, & Prins, 2015). Severe impairment of executive functions in ADHD-C was also suggested in other studies(NIGG, BLASKEY, HUANG-POLLOCK, & RAPPLEY, 2002; Nikolas & Nigg, 2013). A study on time perception showed that the way ADHD-I individuals experience time differs from that of ADHD-C, and this difference is believed to be related to structural variations in the medial frontal cortex, supporting the notion of significant neurocognitive differences between ADHD subtypes. (Bluschke, Schuster, Roessner, & Beste, 2018). However, despite the reports of differences in cognitive functions between ADHD-I and ADHD-C, there is also evidence indicating similar performance in cognitive abilities for both subtypes (Geurts, Verté, Oosterlaan, Roeyers, & Sergeant, 2005; Skogli, Egeland, Andersen, Hovik, & Øie, 2014).

Inconsistent results are also observed in the structural neuroimaging studies. Although several structural and diffusion tensor imaging (DTI) studies have revealed regional structure differences between ADHD-I and ADHD-C in hippocampal(Al-Amin, Zinchenko, & Geyer, 2018), thalamus(Lei et al., 2014), caudate(Lei et al., 2014; Semrud-Clikeman, Pliszka, Bledsoe, & Lancaster, 2012), and various cortical regions(Ercan et al., 2016; Lei et al., 2014; Semrud-Clikeman et al., 2012; Z.-M. Wu et al., 2022), there are frequent reports suggesting no difference in local and global white matter and gray matter between ADHD subtypes(S Carmona et al., 2005; Susanna Carmona et al., 2009; Vilgis, Sun, Chen, Silk, & Vance, 2016). Furthermore, these later studies revealed a few significant differences between ADHD subtypes and normal developing control groups, implying only slight variations in brain structure between individuals with and without ADHD (Faraone et al., 2021).

The inconsistent results in the studies may be attributed to a number of factors, including age, gender, type of medication, and statistical methods used. However, it has been proposed that ADHD is not linked to a specific structural brain issue, but rather, it is more likely related to impairments in functional neural systems(Castellanos & Proal, 2012; Cortese et al., 2012) which can result in a broad range of behavioral and cognitive symptoms. This idea has been supported by almost congruent functional connectivity studies that have shown impairments in the connections between brain regions or deficits in the topology of functional brain network(Cortese et al., 2012; Jacqueline Fifi Saad, Griffiths, & Korgaonkar, 2020; Sutcubasi et al., 2020). Particularly, examinations of neural systems through resting state brain networks have shown changed functional connections in the entire brain network (Amir Hossein Ghaderi et al., 2017; Iravani, Arshamian, Fransson, & Kaboodvand, 2021), default mode network (DMN)(Fair et al., 2013; Uddin et al., 2008), salience network (SN)(Cai, Griffiths, Korgaonkar, Williams, & Menon, 2021), cognitive control network (CCN)(Castellanos et al., 2008), sensorimotor network (SMN)(Kucyi, Hove, Biederman, Van Dijk, & Valera, 2015), and visual network (VN)(Hale et al., 2014; Kucyi et al., 2015) in ADHD. These resting state networks refer to the intrinsic connectivity networks within the brain that are active even in the absence of an externally prompted task and can be significantly altered in various cognitive disorders, leading to changes in brain activity and connectivity. Specifically, some of the resting state brain network studies in ADHD have investigated differences in functional connectivity patterns between ADHD subtypes(Fair et al., 2013; Amir Hossein Ghaderi et al., 2017; Iravani et al., 2021) using functional magnetic resonance imaging (fMRI) and electroencephalography (EEG).

Among different electrophysiological and neuroimaging, EEG is suggested as a reliable complement approach to discriminate children with ADHD. Various features of quantitative EEG such as increased theta/beta ratio(Arns, Conners, & Kraemer, 2013) and other power related biomarkers(Jouzizadeh, Khanbabaie, & Ghaderi, 2020; Kiiski et al., 2020; Koehler et al., 2009; Rodríguez-Martínez et al., 2020; Rommel et al., 2017) have been highlighted as EEG features in ADHD. However, to date the ADHD related EEG features have been mostly observed over specific EEG electrodes or brain regions which can not explain the higher scale systems/networks alterations in ADHD. We previously performed a whole functional brain network analysis and showed significant differences in the brain network integration and segregation between three groups of ADHD-I, ADHD-C, and normal developing control(Amir Hossein Ghaderi et al., 2017). However, a more precise system-based analysis can clarify the possible abnormalities of functional brain networks that are possibly involved in characterization of ADHD subtypes.

To this aim, we used resting state source localized EEG which can provide high spatial and temporal resolution of neural activities(Roberto D Pascual-Marqui et al., 1999; Thatcher, North, & Biver, 2012). This type of data is useful to evaluate functional connectivity measures and functional brain networks in health and disease(A. Ghaderi, Niemeier, & Crawford, 2022b; Amir H. Ghaderi, Andevari, & Sowman, 2018; Jouzizadeh, Ghaderi, Cheraghmakani, Baghbanian, & Khanbabaie, 2021). Further, using graph theoretical analysis, we are able to evaluate properties of modular neural systems (subnetworks) which are constructed by regions communicating with each other in a synchronized shape(Sporns & Betzel, 2016). One commonly used method to assess these modular subnetworks is the Newman modularity algorithm (Newman, 2006). In the current study, we used this algorithm to compare modularity of three well-studied control (default mode, salience, cognitive control), a sensorimotor, and visual networks between ADHD subtypes and normal developing control group. This analysis aims to uncover any alterations in the connectivity within the modular systems that are thought to play a role in ADHD. Furthermore, we also looked into the inter-connectivity between the functional brain networks, which is crucial for information integration in the brain. To do this, we evaluated the average edge betweenness centrality(Jalili, 2016) between the different functional brain networks. This analysis can uncover potential alterations in inter-modular functional connectivity in individuals with ADHD.

To the best of our knowledge, there is no pervious study to investigate the modularity of subnetworks and communication between them in ADHD. However, according to previous studies which have reported functional connectivity abnormalities in these subnetworks in ADHD(Hale et al., 2014; Kucyi et al., 2015; Jacqueline Fifi Saad et al., 2020), we expect that ADHD affect on this inter/intra-modular functional connectivity which can be different between subtypes of ADHD. Understanding of these alterations, can provide more information regarding the neural basis of ADHD at the level of neural systems and networks.

## 2) Method

### 2-1) Participants

we reanalyzed a dataset includes EEG and behavioural data from 35 children (age between 7 and 11 years). Based on the clinical assessments (the child behavior checklist(Tehrani-Doost, Shahrivar, Pakbaz, Rezaie, & Ahmadi, 2011) and the Swanson, Nolan, and Pelham IV (SNAP IV) questionnaire(Hooshyari, Mohammadi, & Delavar, 2008)) performed by child psychiatric, children with ADHD were divided into two ADHD-I (N=10; 4 girls; age: 8.60±1.42 y), and ADHD-C (N=12, 4 girls; age: 8.42±1.78 y) groups. The mean SNAP-IV scores for the ADHD-I group in items 1-9 (related to inattentively) were 2.31 (SD=0.18) and for the ADHD-C group were 1.78 (SD=0.26). The average of SNAP-IV scores for the ADHD-I group in items 10-18 (related to hyperactivity/impulsivity) were 0.57 (SD=0.23) and for the ADHD-C group were 1.86 (SD=0.29). This dataset also contains normal-developing control group (N=13, 5 girls; age: 8.92±1.38 y). Importantly, none of the children in this study had ever taken medication to treat the ADHD or had ever been treated with alternative approaches like neurofeedback which is the unique aspect of this dataset. The original study was approved by the ethical committee at University of Tabriz, Tabriz, Iran, and all methods were accomplished in accordance with the Declaration of Helsinki.

### 2-2) EEG acquisition and preprocessing

EEG acquisition was performed using a Mitsar® amplifier (WinEEG software) with 21 channels in 10/20 standard system (linked ear reference and AFZ as ground). Impedances between electrodes and scalp kept under 10 kΩ and EEG sampling rate was 250 Hz. Under supervision of an EEG technician, EEG were recorded in an isolated quite room for 5 minutes in the resting state eyes open condition. During the EEG recording session, participants were seated in a comfortable fixed chair, facing a blank wall with a plus sign, which served as the fixation point for their gaze. To control the visual input and eye movements, participants were instructed to maintain their gaze on the fixation point throughout the recording. However, they were also instructed to perform appropriate blinks to avoid gaze fixation (which can cause to visual fatigue or the interruption of the flow of visual information). To remove the effects of power source on the EEG signals, we have applied an online 50 Hz notch filter.

Preprocessing was performed by EEGLAB toolbox (https://sccn.ucsd.edu/eeglab/index.php) in MATLAB 2019b. For the EEG preprocessing, first, we applied a basic FIR bandpass filter between 0.1Hz and 40Hz. Then, we segmented EEG signals to 4-seconds length epochs and all segmented epochs were visually inspected. Epochs with artifacts (including eyes-movement, eye blinks, body movement, and muscle activity) were removed. This step was initially performed manually by an expert, and subsequently, we used the *Clean Rawdata* tool in EEGlab to remove any remaining epochs with artifacts after the initial expert artifact rejection. Among all 35 EEGs, the minimum number of artifact-free epochs was 42 which was enough for further analysis. Then, we randomly selected 42 artifact-free epochs for each participant to unify number of epochs between three groups. According to the good quality of EEGs and enough artifact-free epochs, we did not apply artifact correction algorithms like independent component analysis (ICA). Finally, we extracted this data to perform source localization in the LORETA software.

### 2-3) EEG source localization and connectivity analysis by LORETA algorithm

For each individual, forty-two artifact-free EEG segments were analyzed by the LORETA software (version 20190617; http://www.uzh.ch/keyinst/loreta). We used standard LORETA (sLORETA) algorithm to estimate current densities in EEG sources in the frequency domain. This algorithm estimates the source localized current densities in 2,394 gray matter cortical voxels based on the linear smoothness solutions for the EEG inverse problem(R. D. Pascual-Marqui, 2002). The linear smoothness solution in LORETA algorithm is inspired the neurophysiological behaviour of neighboring neuronal populations that show local coupling and are linearly correlated(Roberto D Pascual-Marqui et al., 1999). Recent studies suggest that the accuracy of source localized current density estimation in sLORETA algorithm is acceptable even with limited number of EEG electrodes (Asadzadeh, Yousefi Rezaii, Beheshti, Delpak, & Meshgini, 2020; Cannon et al., 2012; Mégevand & Seeck, 2018; Thatcher et al., 2012). After estimating the localized current densities in cortical voxels, current densities were averaged over 84 regions of interest (ROIs), based on the Talairach coordinates of Brodmann areas (BA) which is available in connectivity-1 toolbox in the LORETA software. The ROIs were cortical BAs including BAs 1 to 11, 13, 17 to 25, and 27 to 47 in both hemispheres.

Then, using connectivity-1 toolbox in the LORETA software, we calculated functional connectivity between all ROIs. To avoid the volume conduction effects on functional connectivity analysis we selected *lagged coherence* as the measure of functional connectivity which contains imaginary part of *coherence* and is a robust measure against volume conduction. Volume conduction refers to the fact that the activity of a source can be picked up by all the EEG electrodes, with zero or near-zero phase delays, which can artificially increase coherence. Lagged coherence excludes the non-lagged part of coherence, which comprises the effects of volume conduction and non-recorded sources that simultaneously drive recorded sources. By excluding the zero-lag contribution, lagged coherence minimizes the effects of volume conduction artifacts and provides a measure of physiological information that is less affected by them(Amir H. Ghaderi et al., 2018; Roberto D. Pascual-Marqui et al., 2011). Functional connectivity analysis was performed in eight EEG bands (sub-bands) including: delta (1-4Hz), theta (4.5-7.5Hz), alpha1(8-10Hz), alpha2 (10.5-12.5Hz), beta1 (13-15.5Hz), beta2 (16-20.5Hz), beta3 (21-30Hz).

These sub-bands of EEG have been frequently employed in cognitive research and have been reported to be implicated in various cognitive dysfunctions across different disorders. The division of alpha into two sub-bands and beta into three sub-bands may be associated with different aspects of attention and sensory processing, and movements. For example, the lower sub-band of the alpha (alpha1) is often associated with relaxed, unfocused attention, while the alpha2 sub-band is often associated more active cognitive processes, including attention, working memory, mental coordination, and cognitive and memory performance (Klimesch, 1999; Vogt et al, 1998). The low beta sub-band (beta1) is often associated with focused, active attention, and movement imagination while the upper beta sub-bands (beta2, and beta3) are associated with high levels of cognitive engagement and processing (Sarnthein et al, 2006; Müller-Putz and Kaiser 2014). The sub-band frequency analysis was performed by the Fourier algorithm in the LORETA software, and a Hann window was employed as a preprocessing step to enhance the accuracy of the analysis. The Hann window is a widely used technique for tapering data before Fourier analysis, which helps reduce spectral leakage and improve the precision of frequency domain analysis. Finally, connectivity values between all RIOs were arranged as adjacency matrices for each frequency band and each participant. In adjacency matrices, rows/columns are associated with ROIs and the values in the matrix arrays represent functional connectivity values between ROIs corresponding to rows and columns.

### 2-4) Modularity and communication analysis

To evaluate modularity of subnetworks we implemented Newman modularity algorithm (Newman, 2006) in MATLAB 2019b (open access code for modularity analysis is available at https://github.com/AHGhaderi/Amir-Hossein-Ghaderi/commit/df636b2105e578a8969ffa88856e54f3267a40a7). Steps in this algorithm are:

1. Selection of a group of nodes (i.e., brain regions) as a subnetwork.
2. Calculation of the connectivity average between the selected nodes.
3. Random shuffling of connections between nodes in whole network.
4. Again, calculation of the connectivity average between the selected nodes (after randomization).
5. Dividing the calculated values in step 2 (connectivity average in the original subnetwork) by the calculated values in the step 4 (connectivity average in the randomized network).

If the result in step 5 is higher than 1, then one can argue that the connectivity pattern within the group of selected nodes (subnetwork) is more specific than a random connectivity pattern. Therefore, group of selected nodes are connected to each other in a modular way and generates a modular subnetwork.

Here, we constructed five functional subnetworks by selecting the related nodes (Brodmann areas) according to the existing literature: 1) default mode network (DMN)(Greicius, Supekar, Menon, & Dougherty, 2009; Raichle et al., 2001; Smallwood et al., 2021) was constructed by choosing bilateral BAs 7, 9, 10, 23, 31, 39, 46, and 47 which consists of precuneus, superior parietal lobule, dorsolateral prefrontal cortex, frontopolar prefrontal cortex, posterior cingulate cortex, angular gyrus, and orbital part of inferior frontal gyrus (figure 1-a). 2) central control network (CCN) (Gitelman et al., 1999; Menon & D’Esposito, 2022; Niendam et al., 2012) was generated by selecting bilateral BAs 6, 7, 8, 9, 24, 40, 44, 45, and 46 which includes premotor cortex, precuneus, dorsolateral prefrontal cortex, lateral premotor cortex (frontal eye field), anterior cingulate cortex, supramarginal cortex, and inferior frontal gyrus (figure 1-b). 3) salience network (SN)(Seeley, 2019; Seeley et al., 2007) was made by considering bilateral BAs 13, 24, 25, and 32 which consists insular cortex, anterior cingulate cortex, and subgenual anterior cingulate cortex (figure 1-c). 4) visual network (VN)(A. Ghaderi et al., 2022b; Wang et al., 2008) was constructed by involving bilateral BAs 8, 17, 18, 19, 21, and 39 that includes frontal eye filed, visual cortices (primary, secondary, and associative; V1-V5), middle temporal gyrus, and inferior parietal cortex (figure 1-d). We calculated the modularity of these subnetworks in all investigated frequency bands. 5) sensorimotor network (SMN)(Zhou et al., 2014) was generated by selecting bilateral BAs 1, 2, 3, 4, and 6 which contains primary somatosensory cortex, primary motor cortex, and premotor cortex (figure 1-e).

**Figure 1:**
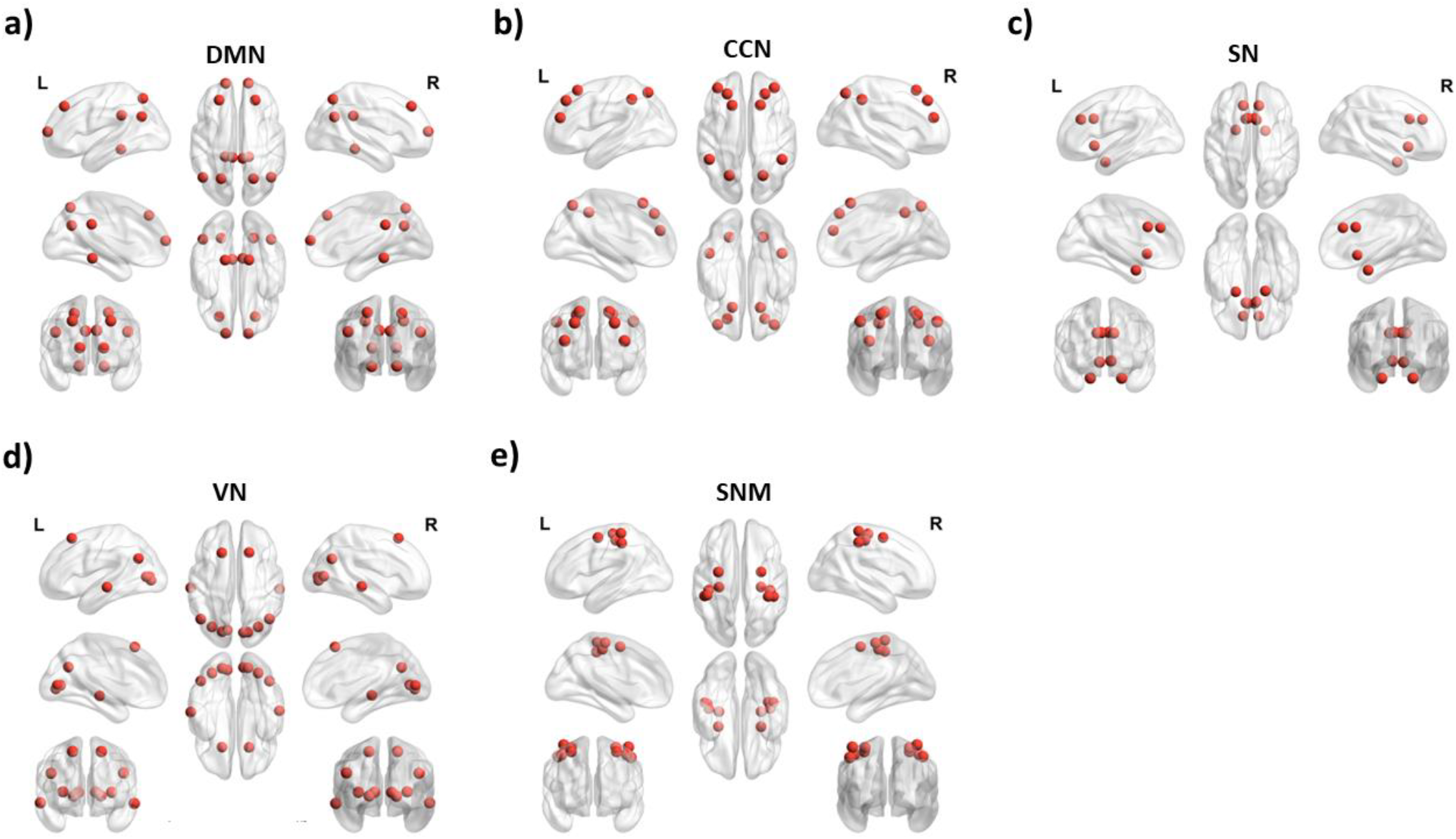
Morphology of functional brain networks. a) The default mode network (DMN) is comprised of bilateral Brodmann areas 7, 9, 10, 23, 31, 39, 46, and 47, and includes the precuneus, superior parietal lobule, dorsolateral prefrontal cortex, frontopolar prefrontal cortex, posterior cingulate cortex, angular gyrus, and orbital part of inferior frontal gyrus. b) The central control network (CCN) is made up of bilateral Brodmann areas 6, 7, 8, 9, 24, 40, 44, 45, and 46 and encompasses the premotor cortex, precuneus, dorsolateral prefrontal cortex, lateral premotor cortex (frontal eye field), anterior cingulate cortex, supramarginal cortex, and inferior frontal gyrus. c) The salience network (SN) consists of bilateral Brodmann areas 13, 24, 25, and 32, which include the insular cortex, anterior cingulate cortex, and subgenual anterior cingulate cortex. d) The visual network (VN) is formed by bilateral Brodmann areas 8, 17, 18, 19, 21, and 39, which encompasses the frontal eye field, visual cortices (primary, secondary, and associative; V1-V5), middle temporal gyrus, and inferior parietal cortex. e) The sensorimotor network (SMN) is made up of bilateral Brodmann areas 1, 2, 3, 4, and 6, and includes the primary somatosensory cortex, primary motor cortex, and premotor cortex.

Then, to evaluate communication between two subnetworks, we calculated edge betweenness centrality (which is equal to number of shortest paths crossing the edges between pair of nodes) between all nodes in two subnetworks (figure 1-f). For example, to evaluate the communication between DMN and SN we calculated edge betweenness centrality between all BAs in DMN and all BAs in SN. Then we averaged these values and higher average edge betweenness centrality between two subnetworks can show higher communication between them.

### 2-5) Data analysis

As there was no prior research on which EEG frequency band could reveal the modular activity of different subnetworks, we conducted an initial investigation into the modularity of all EEG frequency bands in the normal-developing group. Afterwards, we selected the band with the highest modularity for each subnetwork for further analysis. This method prevented us from having to make multiple comparisons and preserved the significance of our results after applying a correction for multiple comparison. In the next step, we compared modularity values (in selected EEG bands) between three groups using permutation t-tests. To correct the multiple comparisons error induced by all comparisons between groups/subnetworks (3 groups ×5 subnetworks= 15 comparisons), a false discovery rate (FDR) analysis was performed, and corrected p-values were reported. Then, to evaluate the communication between subnetwork, we calculated the average of edge betweenness between subnetworks. However, again to avoid many comparisons, we only considered the communications between subnetworks that showed significant modularity deference between groups in the previous step. For the statistical evaluation, we used separate nonparametric permutation t-tests and FDR analysis to correct p-values for 12 comparisons (3 groups and 4 subnetwork communications).

The statistical analysis was accomplished in MATLAB 2019b. For permutation t-test we used a self developed function(A. Ghaderi, Niemeier, & Crawford, 2022a; A. Ghaderi et al., 2022b; Amir Hossein Ghaderi et al., 2018) (open access code for nonparametric permutation analysis is available at https://github.com/AHGhaderi/Amir-Hossein-Ghaderi/commit/4f2c8f731d82707cc06dc20e04fd3ca01b9e8e02) and FDR analysis was performed by *fdr_bh* function which is available at: https://www.mathworks.com/matlabcentral/fileexchange/27418-fdr_bh.

## Results

To reveal which EEG frequency bands exhibited the highest modularity in the resting state brain networks, we first calculated the modularises of different brain networks in different frequency bands in the control group (figure 2-a for VN). The band with higher average value of modularity for each network was selected for statistical analysis. We observed higher modularity of DMN in alpha1 band, SN in delta band, CCN in beta2 band, SMN in beta3, and VN in alpha1. Then, we compared the modularity of these functional brain network in the selected between three groups (i.e., control, ADHD-I, ADHD-C). FDR correction revealed significant differences in modularity of visual network (in the alpha1 band) between the control group and the ADHD-I (t=2.71, q=0.019) and the ADHD-C (t=3.14, q=0.010) groups (figure 2-b). No significant difference was observed between groups in modularity of DMN, SN, CCN, and SMN.

**Figure 2.**
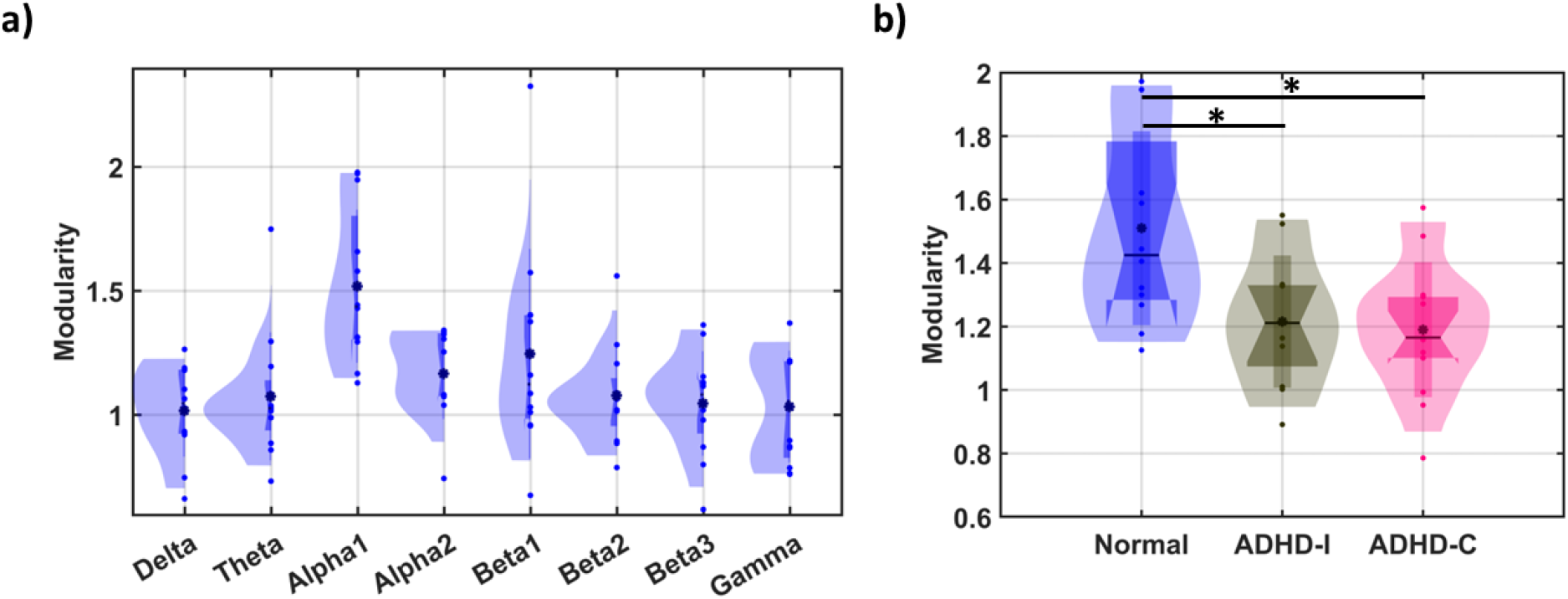
a) Modularity of visual network (VN) in different bands. Values higher than 1 shows modular architecture in subnetwork. b) comparison of VN modularity between three groups in the alpha band. Blue indicates the control group, gray is assigned to the ADHD-I group, and pink is dedicated to ADHD-C. * indicates significant difference alpha<0.05.

Since we observed a significant VN modularity alteration in ADHD subtypes (in the alpha1 band), we then investigated the functional communications between VN and other resting state functional networks (DMN, CCN, SN, SMN). We calculated edge betweenness centrality (as a measure of functional communication between brain regions) between all regions of VN and regions of other networks. Then we averaged the edge betweenness centrality values between all regions for these network pairs: VN-DMN, VN-CCN, VN-SN, and VN-SMN. These values then compared between groups. Figure 3 exhibits the average edge betweenness centralities between network pairs for three groups. In the alpha1 band, for all network pairs the control group showed significant higher average edge between centrality in comparison two ADHD subtypes (statistical details in the figure caption). No significant difference was observed between two ADHD subtypes.

**Figure 3.**
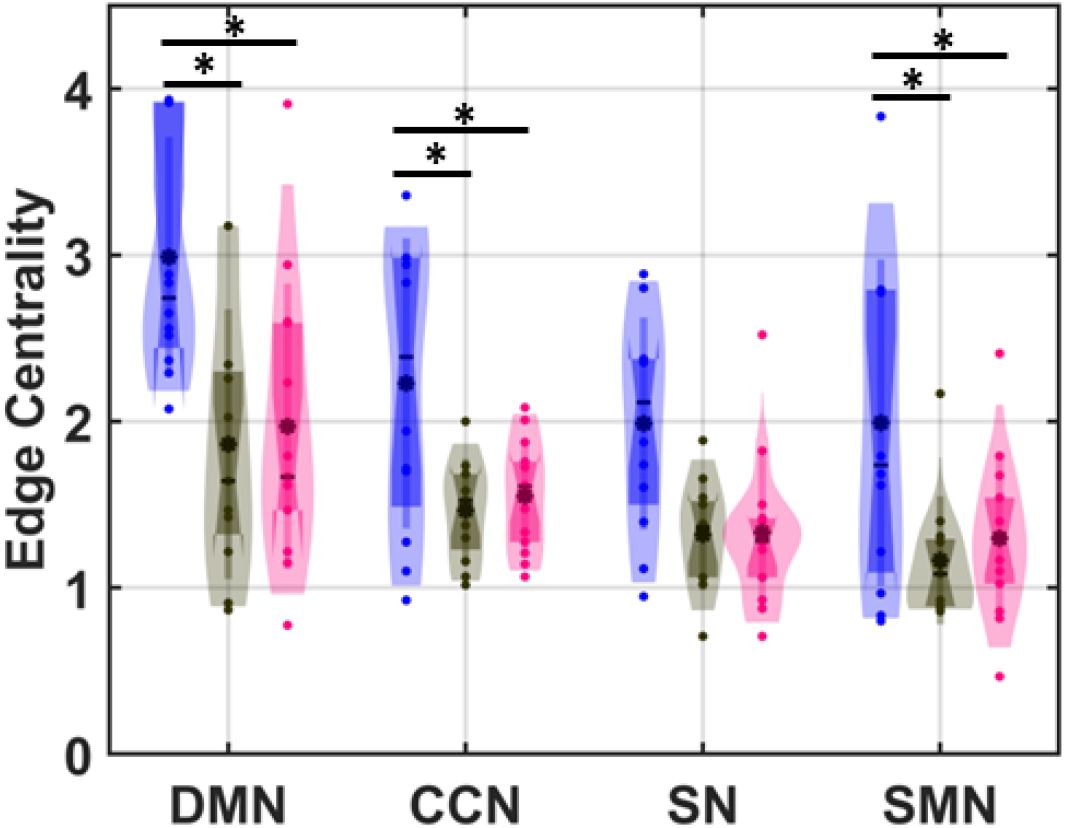
Average edge betweenness centrality between visual network (VN) and four functional brain networks: default mode network (DMN), central control network (CCN), salience network (SN), and sensory-motor network (SMN). Blue indicates the control group, gray is assigned to the ADHD-I group, and pink is dedicated to ADHD-C. According to the FDR analysis, significant differences were observed between the control group (blue) and ADHD-I (gray) for communication between VN and DMN (t=3.52, q=0.023), VN and CCN (t=2.74, q=0.030), VN and SN (t=3.10, q=0.023), and VN and SMN (t=2.61, q=0.030). Also, we obtained significant differences between the control group (blue) and ADHD-C (pink) for communication between VN and DMN (t=3.20, q=0.023), VN and CCN (t=2.60, q=0.030), VN and SN (t=2.96, q=0.023), and VN and SMN (t=2.26, q=0.048). * indicates significant difference alpha<0.05.

## 4) Discussion

Briefly, the results of this study showed that 1) different functional brain networks exhibit modularity in specific frequency bands, 2) ADHD groups showed significant differences in modularity of visual network, 3) the communication between visual network and other functional brain networks (except salience network) was significantly different for ADHD groups in comparison to normal, and 4) no significant differences in modularity of brain networks and communications among them were observed between two groups of ADHD-subtypes. In the section we discuss these results in detail.

### 4-1) Frequency dependent modularity and functional brain networks

Our results showed that each functional brain network exhibits modular connectivity in the specific frequency bands. This result agrees with previous findings suggesting different synchronization states in the brain are associated with various brain functions(Cohen, 2017; Kropotov, 2009).

#### DMN)

According to our results, maximum modularity of DMN was occurred within the alpha band. The modular activity of DMN regions in the alpha band can be justified by previous findings which have suggested increase of alpha synchronization in resting state during imagination and internal attention(Klimesch, 2012; Klimesch, Sauseng, & Hanslmayr, 2007). This result is consistent with previous studies that have used simultaneous EEG/fMRI recordings and showed dominant alpha activity in EEG, when regions of DMN are activated during the resting state condition(Bowman et al., 2017; Mayhew, Ostwald, Porcaro, & Bagshaw, 2013). Further, in consistent with other functional connectivity studies, our result (modular functional connectivity pattern between regions of DMN) agrees with a previous studies that reported long-range synchronization in the alpha band between DMN regions(Jann et al., 2009; Knyazev, Slobodskoj-Plusnin, Bocharov, & Pylkova, 2011). The consistency between our results (that originally obtained by 19 channel EEG recording) and previous simultaneous EEG-fMRI studies exhibits that our proposed algorithm is a valid algorithm and can reveal results which were achieved from more complex setups using simultaneous EEG and fMRI.

#### CCN)

According to our results, functional connectivity in Beta2 exhibited highest modularity of CCN. CCN consists of frontoparietal, and cingulate regions. Functionally, the CCN involves in many cognitive functions including executive functions control(Niendam et al., 2012), visual search(Cole & Schneider, 2007), and actions planning under uncertainty situation(T. Wu et al., 2018; T. Wu, Schulz, & Fan, 2021). Then, the modular activity of this network in the beta band is not surprising because beta activity is thought to be in relation with cognitive and motor processes such as attention, working memory, perception, and motor planning(Fries, 2015; A. Ghaderi et al., 2022b; Amir H. Ghaderi et al., 2018; Tzagarakis, Ince, Leuthold, & Pellizzer, 2010). Increased beta synchronization has been linked to states of arousal and active cognitive processing(Fries, 2015).

#### SN)

SN is another functional brain network (consisting of insula and anterior cingulate cortex regions) with an intrinsic connectivity in the resting state condition(Cai et al., 2021; Menon & Uddin, 2010; Seeley, 2019; Seeley et al., 2007; Zhang et al., 2015). This network is involved in attention allocation to the cognitive or emotional salience stimuli(Goulden et al., 2014; Menon & Uddin, 2010; Seeley, 2019). We observed that modularity of SN is occurred in the delta band. This result agrees with previous studies that suggested detection of motivational salience is accompanied with increase of delta synchronization(Gray, 1999; Knyazev, 2012).

#### SMN)

SMN contains primary motor and sensory cortices and functionally integrates motor and sensory signals. Our result showed that the most dominant modular activity of SMN was occurred in the beta3 band. Previous studies showed several brain waves are involved in activity of SMN, but the most important sensorimotor rhythms are in the range of alpha and beta bands (8-30 Hz). Our results also confirmed the modular activity of SMN in all the alpha and beta ranges, however the strongest modularity was observed in the beta3 band. The prominence of beta wave activity in the sensorimotor network can be attributed to the fact that beta oscillations are particularly strong in the sensorimotor cortex, whereas alpha activity is more widespread and is dominant in larger network including the parietal and sensorimotor regions(Pfurtscheller & Lopes da Silva, 1999; Vukelić et al., 2014).

#### VN)

Visual sensory and visual motor control signals are combined in VN. Many studies have demonstrated that alpha and beta are dominant brain oscillation in the VN regions with more emphasis on alpha. Our result which showed modular activity of VN in the alpha band is not surprising and agrees with previous findings.

### 4-2) ADHD characterized by visual network modularity impairments but preserved sensorimotor and control networks modularity

Interestingly, our results showed that modularity in the resting state visual network is decreased in both ADHD groups in comparison with the normal developing group. Since visual network is responsible for processing and interpreting visual information, reduction of modularity in this network suggests decreased visual processing in children with ADHD. This result agrees with previous findings that explored visual sensory processing impairments in ADHD(Fan, Gau, & Chou, 2014; Ghanizadeh, 2011; McAvinue et al., 2015). These previous studies have reported visual processing impairments during visual tasks, such as processing numbers in the counting Stroop task(Fan et al., 2014). Furthermore, a different lateralization of resting state functional connectivity in visual regions was previously reported in the ADHD group in comparison with normal developing children (Hale et al., 2014). However, to the best of our knowledge, this is the first study that shows impairments in the modularity of the visual network in ADHD during resting state which can be considered as a neural origin for the visual sensory processing deficits in ADHD. It is noteworthy that the decrease in modularity was observed during the resting state, revealing a decline in the intrinsic functional connections between visual-related regions in individuals with ADHD.

Contrary to our expectations, there were no significant differences found in the modularity of other control and sensorimotor networks between the groups. This result suggests that in the subsystem level, ADHD is not related to localized modular connectivity of control or sensorimotor networks. This result may appear to contradict previous findings that suggested functional brain network impairments in the control, or sensorimotor subnetworks(Cai et al., 2021; Castellanos et al., 2008; Kucyi et al., 2015). Notably, this finding differs from the hypothesis that pediatric ADHD is a disorder characterized by decreased connectivity within the core sub-system of the DMN, as suggested by several studies(Fair et al., 2013; Qiu et al., 2011; Sutcubasi et al., 2020; Uddin et al., 2008). The differing results may stem from differences in methodology between our measure of modularity(Newman, 2006) and the average functional connectivity reported in prior studies. Our measure focuses on connectivity within subnetworks compared to the entire network, while prior studies did not take subnetwork connectivity into account. This highlights new insights into the neural effects of ADHD on functional brain networks, suggesting that it does not disrupt modular connectivity in control and sensorimotor brain networks.

Despite unchanged modularity in the control and sensorimotor networks, communication between these networks and the VN was reduced in ADHD (only connectivity between the SN and VN was not different between ADHD and normal developing groups). This is significant as information flow from the VN into the two control networks (CCN and DMN) plays a crucial role in attention, perception, and cognitive control. Hence, we consider this disruption to be a significant neural deficit in ADHD. Further, the reduced communication between the VN and SMN can be seen as the neural basis for disturbed visual-sensory motor integration in ADHD (Ghanizadeh, 2011; McCracken et al., 2019).

Overall, our results uncover a novel functional mechanism for ADHD that can be distilled into three steps. Firstly, lower intrinsic modularity in the VN leads to reduced visual sensory processing. Secondly, disturbed communication between the VN and other networks results in poor sharing of this visual information.

Thirdly, while modularity of control and sensorimotor networks is not impacted by ADHD, the impaired sensory inputs may still impact the function of these networks.

### 4-3) Network modularity differentiates between ADHD subtypes

The study found that the organization of brain activity, as measured by modularity in functional brain networks, was not significantly different between individuals with different subtypes of ADHD. Specifically, individuals with the inattentive (ADHD-I) and combined (ADHD-C) subtypes both showed similar patterns of functional connectivity within and across different brain networks.

This result suggests that there may be a common neural basis for both ADHD subtypes, involving a shared disturbance in the modularity and connectivity of the ventral network. These findings support the current DSM classification of ADHD-I and ADHD-C as two subtypes within the same category, rather than distinct disorders. Despite the differences in symptoms between the two subtypes, the similarities in brain activity organization indicate that both may have a shared underlying neural mechanism. This result contradicts previous studies that suggest ADHD-I is a separate disorder from ADHD-C. It highlights the need for further research to gain a better understanding of the neural mechanisms underlying ADHD and its subtypes.

## 5) Conclusion

In conclusion, the results of this study showed that different functional brain networks exhibit frequency-dependent modularity and that ADHD groups showed significant differences in the modularity of the visual network compared to normal. The communication between the visual network and other functional brain networks, except the salience network, was also found to be significantly different in ADHD groups. These findings suggest impairments in visual processing in children with ADHD and provide insights into the neural mechanisms underlying ADHD. The results also highlight the importance of considering frequency-dependent modularity when studying functional brain networks and their interactions in the context of neurological disorders.

For future research, it is crucial to further explore the underlying causes of these modularity differences in various brain networks and their specific implications for ADHD symptomatology. Investigating the potential role of these findings in the development of targeted interventions or therapies for individuals with ADHD can be a promising avenue. Furthermore, the consideration of frequency-dependent modularity when studying functional brain networks should be integrated into future studies of neurological disorders. This approach could shed light on other conditions and provide a deeper understanding of how different brain networks are affected.

## 6) Limitations

The study was conducted at a single center in Iran and may not generalize to other populations or cultural contexts. Furthermore, the sample size of 35 children is relatively small and may not be representative of a larger population. Nevertheless, this dataset comprised individuals who had not undergone ADHD medication or neuromodulation treatments prior to the study. To address the limited sample size, non-parametric statistical tests were employed to evaluate intergroup disparities, a suitable approach for analyzing datasets with a small number of participants. Also, the EEG source localization and connectivity analysis was performed using LORETA software, which may have limitations in accurately estimating the current densities in EEG sources. However, new studies have shown promising results of LORETA which is comparable with fMRI studies(Asadzadeh et al., 2020; Kuar et al., 2022; A. Ghaderi et al., 2023a; Ghaderi et al., 2023b). Lastly, a limitation of assessing lagged coherence lies in the potential for spurious interactions, which can affect the accuracy of calculations. However, employing source localized EEG helps mitigate this effect, bolstering the robustness of lagged coherence as a connectivity measure(Palva et al., 2018). Nonetheless, future studies could explore alternative functional connectivity measures to further elucidate and possibly mitigate the impact of these spurious interactions, providing additional clarity in our understanding of brain modularity in ADHD.

